# Cellular composition and the interpretation of candidate RNA readouts of trogocytosis in colorectal cancer

**DOI:** 10.64898/2026.10.28.762407

**Authors:** Jinglin Wang, Pan Sun, Lei Xu, Yao Wang, Wei Zhu, Yan Zhang, Yao Tang, Juan Yang

**Author notes:** Corresponding author: Juan Yang, MD, PhD; Department of Geriatrics, The Affiliated Hospital of Guizhou Medical University, Guiyang, China.

## Abstract

**Background:** RNA measurements of genes associated with trogocytosis may reflect cellular composition rather than contact-dependent membrane acquisition. We evaluated an eight-gene candidate module in colorectal cancer (CRC) using expression- and lineage-matched random modules, independent compartment analysis and patient-level inference. A separate rare-transcript definition and cancer-model fitness screens provided complementary analyses.

**Results:** In 62 GSE178341 patients, the module correlated with T-cell fraction (ρ=0.767), but this association was typical of 10,000 matched modules (null median 0.766; one-sided empirical P=0.493; equal-tail two-sided P=0.985). A post hoc closest-10% matching sensitivity gave P=0.571. In 23 independent GSE132465 patients, the tumour-wide correlation was 0.735 (permutation P=0.00060) and the epithelial correlation was 0.376 (P=0.0892). The difference in absolute correlations was 0.359, with a 95% confidence interval of −0.041 to 0.759, leaving attenuation uncertain. Among 43,817 leukocytes, 16 *CEACAM5*-positive candidates arose from four patients; 15 had expression ≤2 transcripts per million. Candidate counts were threshold-sensitive. No survival association was detected for the module in two cohorts. *SCD* fitness dependency reproduced across CRC organoids and cell lines, whereas adjusted *SCD*–*VPS72* and exploratory *ATF3*–*SMYD2* co-dependencies did not. Exploratory *CH25H*, *ATF3* and *FADS2* analyses yielded no Hallmark enrichment meeting the multiple-testing threshold.

**Conclusions:** Matched-module calibration placed the strong association with T-cell abundance within the range expected for comparable immune-lineage genes. Independent compartment analysis was consistent with a contribution from cellular composition, although attenuation remained uncertain. These results identify the comparison conditions needed to interpret the tested RNA readouts. Linking them to membrane transfer requires event-resolved protein measurements.

## Introduction

During trogocytosis, a recipient cell acquires portions of another cell’s membrane and associated proteins through direct contact. Experimental studies have linked this process to host-cell injury in amoebic infection and antigen escape during CAR T-cell treatment [1, 2]. In colorectal cancer (CRC), protein-level and cell-contact experiments have demonstrated acquisition of lymphocyte membrane material by malignant cells, with acquisition-associated populations displaying immune-regulatory proteins including CTLA-4, PD-1, TIM-3 and LAG-3 [3]. These observations make membrane acquisition a relevant experimental phenotype in CRC. They also motivate a distinct genomic question: what can RNA measurements of the corresponding genes reveal about that phenotype?

RNA abundance and membrane-protein acquisition are not interchangeable measurements. A recipient can acquire a surface protein without transcribing its gene, whereas a mixed specimen can contain abundant immune-marker RNA because it contains many immune cells. This distinction is particularly relevant in CRC, where molecular subtypes reflect malignant epithelial programmes together with immune and stromal contributions [4, 5]. Spatial and organoid– fibroblast studies also show that local tissue context shapes epithelial phenotypes [6]. An association between a mixed-marker RNA score and immune-cell abundance therefore needs a reference comparison: is it stronger than the association expected for other genes with similar expression and lineage specificity?

Functional screens offer a complementary view of the genes implicated in membrane-associated biology. Experimental studies implicate *ATF3* and *CH25H* in effector-cell trogocytosis [7]. In colon cancer stem cells, low *CH25H* expression has been linked to greater membrane fluidity and trogocytosis [8]. Such observations motivate lipid-related hypotheses, but cancer-cell fitness screens measure the consequences of gene loss for the screened cells. A dependency identified in a monoculture cannot by itself establish a role in an interaction between donor and recipient cells.

Here, we calibrated an author-assembled, fixed eight-gene RNA module against expression-, detection- and lineage-matched random modules in a public CRC single-cell cohort. We then examined tumour-wide and epithelial pseudobulk scores in an independent cohort and evaluated a separate *CEACAM5*-positive leukocyte definition across patients and detection thresholds.

Molecular context and survival analyses assessed the behaviour of the specified score, while organoid and cell-line screens assessed supporting fitness phenotypes. Together, these analyses assess how cellular composition, compartment choice and detection thresholds shape candidate RNA readouts. The patient datasets lack event-resolved membrane-transfer measurements.

## Materials and methods

### Ethics approval and consent to participate

This study re-analysed publicly available, de-identified datasets and published supplementary tables. No new participants were recruited, and no new human specimens or animal data were collected. No separate ethics approval or formal exemption was obtained for this secondary analysis. At no stage of this study did any author have access to information identifying individual participants or to a key linking study identifiers to personal identities. Ethics approval and consent for the original data collection were the responsibility of the original studies and repositories.

Archived execution records document research access to GSE178341, GSE146771, GSE39582 (curatedCRCData) and TCGA-COAD/READ (cBioPortal PanCancer Atlas) data or their derived analysis inputs on 31 August 2026, and to GSE132465 on 1 September 2026. These dates were reconstructed from project execution records in China Standard Time (UTC+8) and describe documented research use, not necessarily the first download. Access dates for the published organoid tables and DepMap data are reported in the functional-screen methods below.

Clinical trial number: not applicable.

### Consent for publication

Not applicable. No potentially identifiable individual information or images are included.

### Study design and analysis timing

We re-analysed public, de-identified patient datasets and published cancer-model screens. The independent unit was the patient, organoid model or cell line, as appropriate (Table 1). Cell counts were not treated as independent patient replicates. The eight-gene module and the original thresholds were fixed before their corresponding calibration or replication analyses; this freezing followed initial exploratory work and was not a prospective registration. *CH25H*, *ATF3* and *FADS2* comparisons and the *ATF3*–*SMYD2* replication check were added in response to peer review and are explicitly exploratory. Candidate results were available before the independent implementation checks. No new wet-laboratory experiments were performed.

**Table 1.** Public data layers and analysis units.

| **Data layer** | **Analytic size** | **Independent unit** | **Role** |
|---|---|---|---|
| GSE178341 | 257251 tumour cells; 62 patients | Patient | Matched-null calibration and contextual associations |
| GSE132465 | 47285 tumour cells; 23 patients | Patient | Independent compartment analysis |
| GSE146771 | 43817 leukocytes; 8 paired patients | Patient for tissue inference | Sparse RNA candidate sensitivity |
| GSE39582 | 519 univariable / 518 adjusted RFS cases; 145 events | Patient | Relapse-free survival association |
| TCGA-COAD/READ | 566 / 554 OS cases; 115 / 110 deaths | Patient | Overall-survival association |
| CRC organoid screen | 85 models; 16562 complete genes | Organoid model | Supporting functional and co-dependency analyses |
| DepMap Public 26Q1 | 63 CRC and 1145 other lines | Cell line | Functional context and association checks |

RFS, relapse-free survival; OS, overall survival. Numbers denote analytic sets, not the full source-study recruitment.

A trogocytosis event was understood as contact-dependent acquisition of donor membrane material or membrane-associated protein by a recipient cell. RNA scores and rare transcripts were treated as candidate readouts, not event labels. Fitness loss, molecular association, clinical association and direct transfer evidence were kept separate throughout interpretation.

### Fixed module and exploratory gene panel

The RNA module comprised *CD4*, *PTPRC*, *CTLA4*, *PDCD1*, *HAVCR2*, *VSIR*, *LAG3* and *CD38*. Its score was the arithmetic mean of gene-wise standardised expression within each cohort and, for compartment analyses, within each compartment. It was not optimised against survival. This author-assembled combination contains lineage markers alongside genes encoding proteins reported in transfer experiments; it is not an externally validated trogocytosis signature.

We assembled a heterogeneous, literature-informed panel of 39 genes for exploratory analysis. The panel included antigens and regulatory molecules investigated in experimental studies of trogocytosis, immune-interface and phagocytic-checkpoint molecules, and candidates reported in exploratory transcriptomic studies. S1 Table provides gene-level references, inclusion rationales, experimental contexts and evidence categories, distinguishing protein transfer or its regulation from related biological context and transcriptomic association; the readable annotation and its source bibliography are in Table A of S1 Appendix. The panel was author-assembled rather than systematically exhaustive and was used to examine functional dependencies and candidate RNA readouts, not as a validated gene set for quantifying trogocytosis. Five genes (*CD24*, *FCGR1A*, *FCGR2B*, *FCGR3A* and *SIRPA*) were absent from the organoid matrix, leaving 34. *FADS2*, encoding fatty acid desaturase 2, was an additional exploratory comparator outside this panel.

### GSE178341 patient profiles and matched random modules

The complete GSE178341 H5 matrix contained 370,115 cells with author-provided cell and specimen annotations [9]. Tumour samples with mismatch-repair annotation contained 257,251 cells from 62 patients (34 MMR deficient [MMRd] and 28 MMR proficient [MMRp]). The patient identifier (PID), not the specimen-level PatientTypeID, was used as the clinical analysis unit.

Patient pseudobulk profiles were generated for all tumour cells and separately for the epithelial compartment. Consensus molecular subtype (CMS) was assigned from epithelial pseudobulk using the official CMScaller template [10]. Raw calls comprised CMS1 n=17, CMS2 n=23, CMS3 n=14 and CMS4 n=8. A sensitivity analysis retained only calls passing a false discovery rate (FDR)<0.05 (CMS1 n=14, CMS2 n=19, CMS3 n=9, CMS4 n=7 and unclassified n=13). The frozen eight-gene module was computed for whole-tumour and epithelial pseudobulk. T-lineage cells were defined by the author midpoint classes TCD4, TCD8, Tgd and TZBTB16; patient T-cell fraction was their combined count divided by all tumour cells. The author-annotated exhausted-cell fraction was calculated within T-lineage cells from TNK/ILC cluster labels containing *CXCL13* or *PDCD1*. A separate exhaustion-expression score used *PDCD1*, *CTLA4*, *HAVCR2*, *LAG3*, *TIGIT*, *TOX* and *CXCL13*.

Spearman correlations related module scores to transforming growth factor beta (TGF-beta), epithelial–mesenchymal-transition (EMT) and T-cell-fraction measures. Pathway correlations used patient-level bootstrap 95% confidence intervals (CIs) and Benjamini–Hochberg (BH) correction across the four primary unadjusted pathway tests. MMRd–MMRp comparisons included T-cell fraction, author-annotated exhausted T-cell fraction and the exhaustion-expression score; BH correction covered seven group comparisons: these three MMR comparisons and four CMS comparisons crossing whole-tumour/epithelial scores with raw/FDR-filtered CMS calls.

To determine whether the correlation between the frozen module and T-cell fraction exceeded that expected for comparable immune-lineage genes, 10,000 random eight-gene modules were generated in the same 62-patient GSE178341 tumour cohort. Candidate genes were matched to each target gene using three features calculated before module sampling: tumour-wide mean expression, tumour-cell detection rate and log2 TNK/ILC-versus-other-cell pseudobulk specificity. Unique non-mitochondrial and non-ribosomal symbols detected in at least 0.05% of tumour cells, present in at least 10 patient pseudobulks and variable between patients were eligible. The frozen genes and complete 39-gene panel were excluded, leaving 21,904 candidates.

Matching used robustly standardised Euclidean distance, with one gene per target and without replacement. The primary neighbourhood size was k=50; k=25 and k=100 were prespecified sensitivities. Every random module used the same patient-level log2 counts per million (CPM), gene-wise standardisation and mean-score procedure as the frozen module. The prespecified directional test was the empirical upper-tail probability that a matched module correlated at least as strongly as the frozen module. A descriptive two-sided value was defined as twice the smaller empirical tail probability, capped at 1; it was centred on the empirical matched-null distribution rather than on zero. Null intervals and the observed percentile were also reported. A balance audit compared the three frozen-module means with their matched-null distributions. Because the expression mean lay near the upper edge of its null range, an explicitly post hoc sensitivity retained the 10% of primary modules closest to the frozen module across all three standardised module-mean features; it did not replace the prespecified analysis.

The calibration used the complete-gene CPM implementation for both the observed and random modules. This yielded observed ρ=0.767218, compared with 0.766513 in the retained contextual analysis; their patient score ranks were highly concordant (ρ=0.993302). These two implementations round to the same displayed correlation, but their unrounded coefficients and asymptotic P values were not interchanged. The null percentiles and empirical P values refer exclusively to the complete-gene calibration. A large empirical P indicates no unusually strong association relative to the chosen controls; it is not an equivalence test and does not estimate the fraction of the association causally explained by composition.

### Independent compartment analysis in GSE132465

GSE132465 was selected before inspecting module results because it is an independent human CRC atlas with epithelial and immune compartments, more than 10 tumour patients, a public unique molecular identifier (UMI) matrix and author cell annotations [11]. The processed dataset contained 63,689 cells from 23 Korean patients; analysis was restricted to 47,285 tumour cells, including 17,469 epithelial cells and 16,739 T cells. T-cell fraction was defined as author-annotated T cells divided by all tumour cells for each patient. The legacy matrix symbol *C10orf54* was mapped to its current synonym *VSIR*; no other module gene was substituted.

Raw UMI counts and library sizes were aggregated by patient for all tumour cells and separately for author-annotated epithelial cells. The frozen score was the mean of gene-wise standardised log2(CPM+0.25) values. Missing or invariant genes were specified to contribute zero and to be recorded; all eight genes varied in both compartments. Tumour-wide and epithelial scores were correlated with T-cell fraction using Spearman analysis, 10,000 patient bootstraps and 10,000 permutations. The prespecified attenuation statistic was |ρ tumour-wide|−|ρ epithelial|. Strong replication required a positive significant tumour-wide association, attenuation >0.20 and an attenuation bootstrap CI above zero; the same point criteria with a CI crossing zero were labelled compatible replication.

The independent analysis tested whether the pattern was compatible with composition tracking. A significant tumour-wide association and a non-significant epithelial association were not, by themselves, treated as a significant difference between correlations. The bootstrap interval for their absolute-correlation difference was the relevant uncertainty measure.

### Rare *CEACAM5* transcripts in GSE146771

The author-annotated 10x leukocyte atlas GSE146771 contained 43,817 cells: 20,466 from peripheral blood, 12,657 from adjacent normal tissue and 10,694 from tumour [12]. Because the dataset contains RNA but no paired surface-protein or lineage-transfer channel, the analysis sought candidate cells rather than events. The primary definition required *CEACAM5*>1 TPM (transcripts per million) and an epithelial-background score no greater than the 99.5th percentile among peripheral-blood leukocytes. The background score was calculated from 15 non-*CEACAM5* epithelial markers. Tissue rates were accompanied by Wilson 95% CIs.

For the eight patients with paired adjacent-normal and tumour specimens, candidate rates were calculated per patient and tissue. The paired difference was evaluated by an exact sign-flip test. Leave-one-patient-out and threshold-grid analyses varied the peripheral-blood percentile (99.0, 99.5 and 99.9) and *CEACAM5* threshold (>0, >1 and >2 TPM). Cell-level enrichment tests were considered descriptive because the candidate count was low and cells from the same patient are not independent.

This definition examines a different proposed RNA readout from the eight-gene score. No candidate was assigned a validated positive transfer label. The epithelial-background gene set and the complete threshold grid are retained in the original analysis configuration and in the supplied source tables; no threshold was selected to improve a tumour-enrichment result.

### Clinical association analyses

GSE39582 was accessed as the GSE39582_eset object from the Bioconductor curatedCRCData resource and cross-checked against the GEO series description [13]. Gene-level expression values were the frozen-normalised microarray values supplied in that curated object; no cross-cohort renormalisation or outcome-guided feature selection was performed. The object contains 585 clinical records, of which 566 have tumour expression profiles. Its days_to_death and vital_status fields were entirely missing, so overall survival was not reconstructed by mixing endpoints from another release. Relapse-free survival (RFS) models used cases with positive follow-up time and complete event data: 519 cases (145 events) for the univariable model and 518 cases (145 events) after complete-case adjustment for age, categorical stage, mismatch-repair status and adjuvant chemotherapy.

The frozen eight-gene score was standardised to one cohort standard deviation (SD). Cox proportional-hazards models reported hazard ratios (HRs) per SD with 95% CIs. The proportional-hazards assumption was tested using Schoenfeld residuals for the module and globally for each fitted model. Kaplan–Meier curves used a median split only for display; inference used the continuous score.

RNA-seq-v2 RSEM expression and patient-level clinical data were retrieved from the cBioPortal study coadread_tcga_pan_can_atlas_2018 [14]. The API returned 592 patients with expression data. After requiring a positive overall-survival time and non-missing status, 566 patients with 115 deaths formed the univariable analysis. Age- and categorical-stage-adjusted complete-case analysis included 554 patients with 110 deaths. Expression was log2-transformed where appropriate, standardised within gene, and averaged across the same frozen eight genes. For TCGA, gene-wise and composite-score standardisation used sample SDs (ddof=1) in the expression cohort before follow-up filtering; complete-case subsets retained that scaling. The adjusted TCGA model used R survival 3.8-6, Efron handling of ties and categorical stage I–IV. Proportional-hazards diagnostics used cox.zph with the rank time transformation, reporting the module term and the joint GLOBAL test separately. Kaplan–Meier curves again served only as displays. Prespecified sensitivity analyses varied score scaling and complete-case filters without redefining the gene set.

These analyses tested association of the specified score with available outcomes. They did not evaluate the prognostic value of independently measured trogocytosis, and absence of statistical association was not taken as evidence of clinical equivalence.

### Supporting functional screens and exploratory comparisons

Supplementary tables from the tumour-derived organoid biobank [15] were downloaded on 30 August 2026. Of 132 CRC models, 85 had usable CRISPR screens. When a model had been screened with two libraries, the library with the higher study-provided quality-control (QC) order was retained, giving one best-library record per model. Gene-level knockout log fold-changes (LFCs) and study-provided is_depleted calls were extracted. The latter were used for official depletion percentages; no universal LFC threshold was substituted for the study’s criterion. QC covariates included areas under the receiver-operating-characteristic curve (AUC-ROC) and precision–recall curve (AUC-PR).

CMS labels were available for 52 models (CMS1 n=11, CMS2 n=16, CMS3 n=11 and CMS4 n=14). Microsatellite status was joined only after restricting the source annotation to organoid records, preventing whole-genome or tumour records with the same sample name from overwriting organoid status. This yielded 15 microsatellite-unstable (MSI) and 70 microsatellite-stable (MSS) models. Differences in *SCD* dependence across CMS groups were assessed using the Kruskal–Wallis test. MSI–MSS comparisons were performed for all 34 screen-covered panel genes using Mann–Whitney tests followed by BH correction.

DepMap Public 26Q1 CRISPR Chronos gene effects and model annotations were downloaded on 31 August 2026 [16]. Sixty-three CRC cell lines had gene-effect data; 1,145 cell lines from other tumour lineages formed the comparison group. For the 34 panel genes shared with the organoid screen, organoid median LFC was compared with the median CRC Chronos effect using Spearman rank correlation. Because the two platforms differ in scale and in how depletion is defined, the comparison was rank-based and platform-specific rates were displayed separately.

For the original *SCD* and *EGFR* contrasts, CRC-versus-other-lineage Mann–Whitney tests and bootstrap median-difference intervals were retained. Exploratory comparisons of *SCD*, *CH25H*, *ATF3* and *FADS2* used 63 CRC and 1,145 other cell lines, 10,000 bootstrap resamples (seed 42), and BH correction across four lineage tests. These descriptive comparisons do not constitute a comprehensive specificity test of *SCD*. Rates based on study-provided organoid depletion calls were not equated with counts below a numerical LFC or Chronos threshold.

For each of the four anchors, a complete 85-model by 16,562-gene organoid matrix was used. Gene ranks and anchor ranks were residualised against rank-transformed genome-wide median LFC, AUC-ROC and AUC-PR, including an intercept. Pearson correlation of the residual ranks gave the adjusted association. The approximate partial-correlation t statistic used 80 degrees of freedom (85−3−2) consistently in the statistic and its reference distribution. The anchor was excluded before BH correction across 16,561 partners. We report both the single-anchor q values and a sensitivity analysis pooling all four anchor families (66,244 tests; S9 Table). Full tables include negative and non-significant associations.

The original *SCD* preranked analysis used Hallmark 2023.2, gene-set sizes of 15–500, 5,000 gene-set membership permutations and seed 20260831 [17]. BH-adjusted nominal P<0.05 was the threshold. A separate exploratory analysis evaluated *SCD*, *CH25H*, *ATF3* and *FADS2* using the same cached Hallmark 2020-labelled collection for all four anchors. Each complete adjusted-correlation ranking contained 16,561 genes after removing the anchor. Weighted enrichment scores (weight exponent 1) were calibrated against 5,000 gene-set permutations per pathway (seed 42), with same-sign null normalisation and plus-one nominal P values. BH correction was applied to 50 pathways per anchor, with a pooled 200-test sensitivity. These BH-adjusted values are distinct from the pooled-normalised-score FDR estimator used by the Broad GSEA software. The two Hallmark collection versions were not assumed interchangeable; the original *SCD* results and this exploratory extension are reported separately. Supplementary methods and S10 Table provide the input hash, implementation details and complete results.

The organoid *SCD*–*VPS72* and exploratory *ATF3*–*SMYD2* associations were checked in DepMap with a common implementation. Common-essential genes were defined across all 1,208 two-dimensional cell lines using ≥90% availability, median Chronos effect ≤−0.5 and an effect below −0.5 in ≥80% of all lines. Excluding each tested pair left 1,179 genes. Missing values were median-imputed for principal-component construction; effects were standardised across all lines, and the first five components were retained. Within CRC, target ranks were residualised against these components, the per-line genome-wide median and median absolute deviation. Ten thousand CRC-line bootstraps reranked and refitted the target model; 10,000 residual permutations used a plus-one empirical P estimator (seed 20260901). The common-essential basis was held fixed. Replication required the organoid-negative direction, P<0.05 and a bootstrap interval excluding zero. Adjustment mitigates measured shared structure but does not make these observational associations causal.

### Statistical reporting and reproducibility

Effect sizes, confidence intervals, denominators and multiplicity families accompany P values. Tests were two-sided except the declared upper-tail matched-null comparison. Analyses used patient-level aggregation, complete-case clinical models and no clinical-covariate imputation. The gene set and endpoints were retained throughout calibration, compartment analysis and clinical association testing. The exploratory functional analyses used Python 3.12.14, NumPy 2.4.6, pandas 2.3.3 and SciPy 1.18.0. Input hashes, selected model/library pairs, scripts, complete association tables and plotting sources are retained. Algebraic agreement of the partial-correlation and regression t statistics, model uniqueness, self-exclusion and BH family sizes were checked. The original adjusted DepMap *SCD*–*VPS72* estimate was reproduced as an implementation control.

### Use of AI-assisted tools

OpenAI Codex assisted with code checking, language editing and polishing during August– October 2026. These tools were not treated as authors or as scientific sources. Responsibility for the analyses, interpretations and submitted text remains with the authors.

## Results

### The module–T-cell association was typical of matched controls

In GSE178341, the fixed module correlated strongly with T-cell fraction across 62 patients, but its coefficient was typical of the matched random modules (Fig 1A). The complete-gene calibration gave ρ=0.767 and a null median of 0.766, with a central 95% null interval of 0.668– 0.843. The observed score lay at the 50.8th percentile: the one-sided empirical P was 0.493 and the descriptive equal-tail two-sided P was 0.985. The latter describes extremeness relative to the matched distribution, not departure from zero correlation. Alternative neighbourhood sizes gave one-sided P=0.579 for k=25 and P=0.388 for k=100. Complete calibration values for all three neighbourhood sizes and the matching-balance audit are provided in S11 Table; patient-level scores and T-cell fractions are provided in S12 Table.

**Fig 1.**
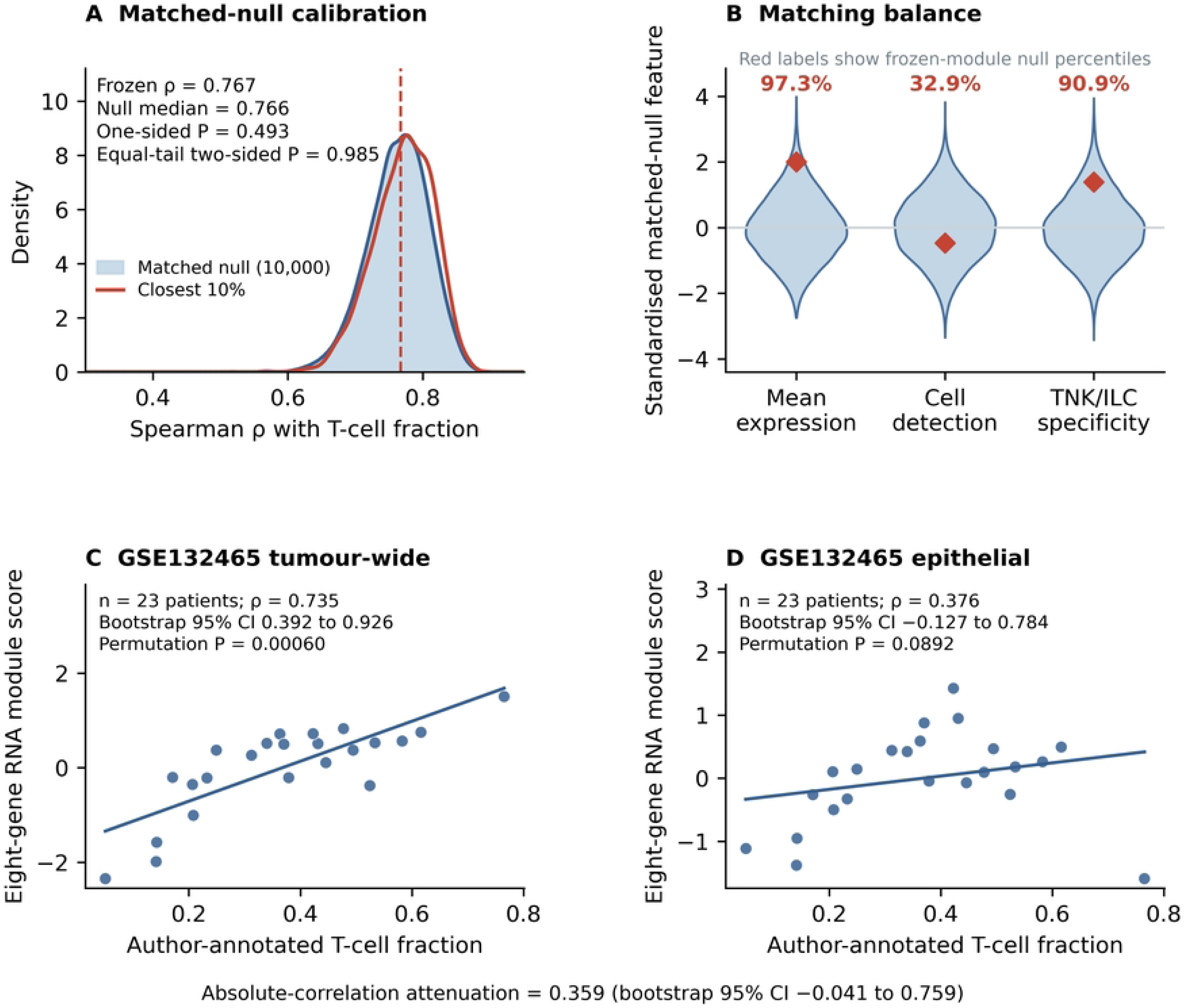
Matched controls and independent compartment analysis of the eight-gene RNA module. (A) Kernel-density displays of 10,000 k=50 matched-module correlations and the post hoc closest-10% subset in 62 GSE178341 patients. The vertical line is the complete-gene observed correlation. P values compare the observed score with the matched distribution, not with zero. (B) Frozen module-mean features (red diamonds) relative to standardised null distributions; red percentages are null percentiles, not proportions of cells. (C,D) Independent GSE132465 patient scores against author-annotated T-cell fraction (23 patients). Lines are linear visual guides; inference uses Spearman correlation, patient bootstraps and permutations. The interval for absolute-correlation attenuation includes zero.

Matching was imperfect at the module level (Fig 1B). Frozen mean expression, detection and TNK/ILC specificity lay at null percentiles of 97.3, 32.9 and 90.9. An explicitly post hoc analysis retaining the 1,000 most closely balanced modules gave a median correlation of 0.774 and P=0.571. Thus, the observed positive correlation did not distinguish the specified module from these immune-lineage-matched controls.

### Tumour-wide and epithelial associations in an independent cohort

GSE132465 contributed 23 patients and 47,285 tumour cells. The tumour-wide score correlated with author-annotated T-cell fraction (ρ=0.735; bootstrap 95% CI 0.392–0.926; permutation P=0.00060; Fig 1C). Restriction to 17,469 epithelial cells yielded a lower estimate (ρ=0.376; 95% CI −0.127 to 0.784; permutation P=0.0892; Fig 1D). All eight module genes varied between patients in both compartments. Cohort-level and patient-level replication results are provided in S13 Table.

The absolute-correlation difference was 0.359, exceeding the prespecified point threshold of 0.20, but its bootstrap 95% CI was −0.041 to 0.759. The analysis therefore met the specified definition of compatible rather than strong replication. It reproduced the tumour-wide association while leaving uncertainty about its attenuation after epithelial restriction. These RNA measurements cannot distinguish biological transcription within epithelial cells from ambient RNA, doublets or annotation leakage.

### *CEACAM5*-positive leukocyte candidates were sparse and threshold-sensitive

The primary filter identified 16 candidates among 43,817 GSE146771 leukocytes, arising from four patients (Fig 2). Fourteen candidates were in adjacent normal tissue, two in tumour and none in peripheral blood. In eight patients with paired normal and tumour specimens, the mean tumour-minus-normal difference was −7.48×10^−4^ per leukocyte (exact sign-flip P=0.125). Leave-one-patient-out tests retained the negative direction, with P=0.125–0.250. These patient-level results do not establish enrichment in either tissue. Complete tissue-level rates and patient-, tissue- and lineage-level counts are provided in S14 Table.

**Fig 2.**
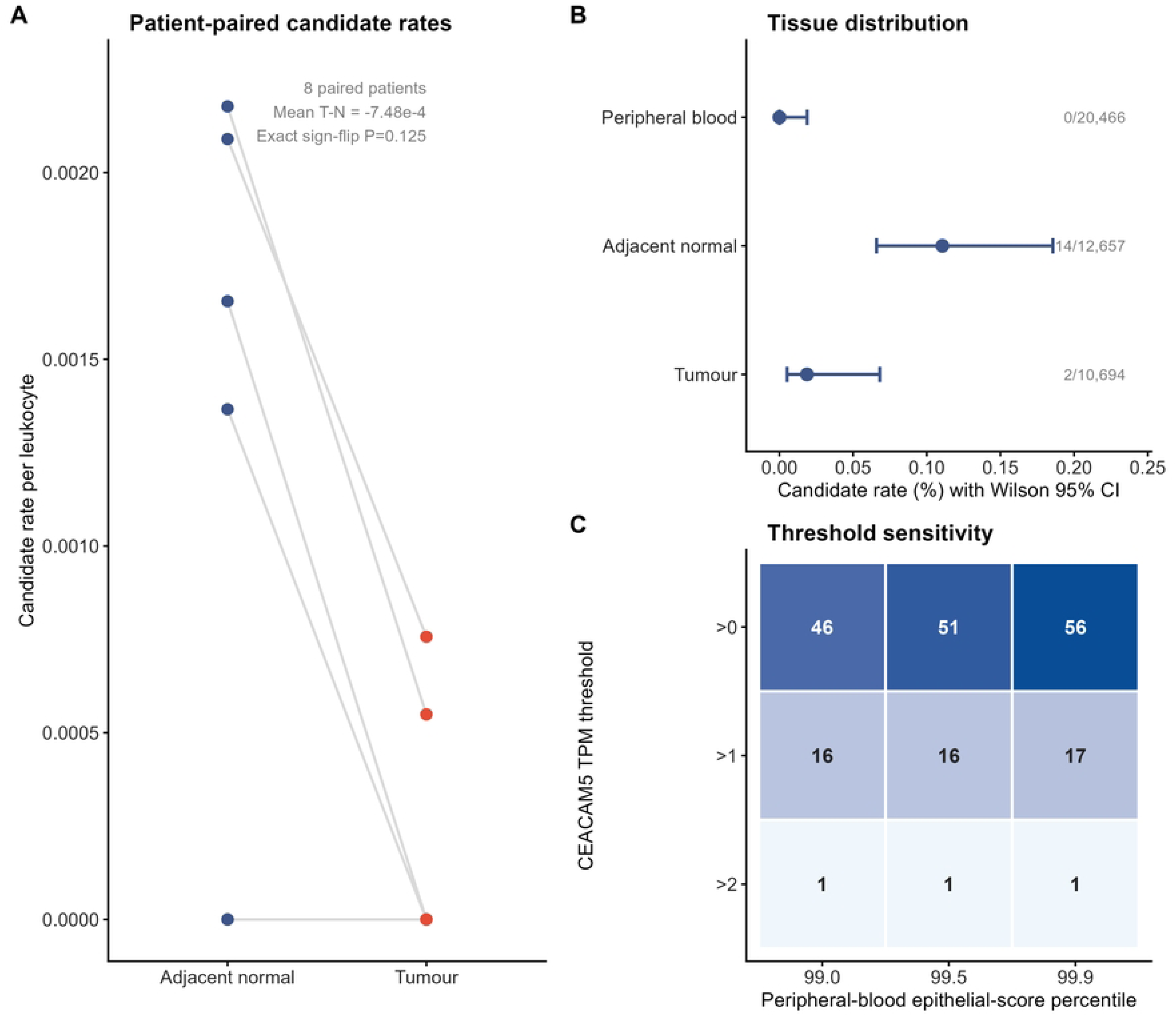
RNA-defined *CEACAM5*-positive leukocyte candidates in GSE146771. (A) Candidate rates in paired adjacent-normal and tumour specimens from eight patients; overlapping zero values are present. (B) Candidate rates by tissue with Wilson 95% CIs and raw counts. (C) Candidate-count sensitivity to *CEACAM5* and epithelial-background thresholds. The primary threshold yielded 16 candidates from four patients; 15 had *CEACAM5* TPM≤2. Candidates are not classified as trogocytosis events.

All 16 primary candidates had zero epithelial-background score, and 15 had CEACAM5 TPM≤2. Across the background-percentile grid, *CEACAM5* thresholds >0, >1 and >2 TPM yielded 46–56, 16–17 and one candidate, respectively. The instability arose principally from low *CEACAM5* expression rather than the background cutoff. Without transfer-positive controls or paired protein measurements, sensitivity and positive predictive value for actual trogocytosis cannot be estimated from these candidates.

### CMS and MMR associations with the RNA module

Aggregation by patient identifier combined 64 tumour specimen records into 62 patient-level units. With raw epithelial-pseudobulk CMScaller labels, neither whole-tumour nor epithelial scores were enriched in CMS4 (P=0.451 and 0.718); filtered calls likewise gave P=0.856 and 0.944 (Fig 3A,B). These subgroup results were not used to redefine the module.

**Fig 3.**
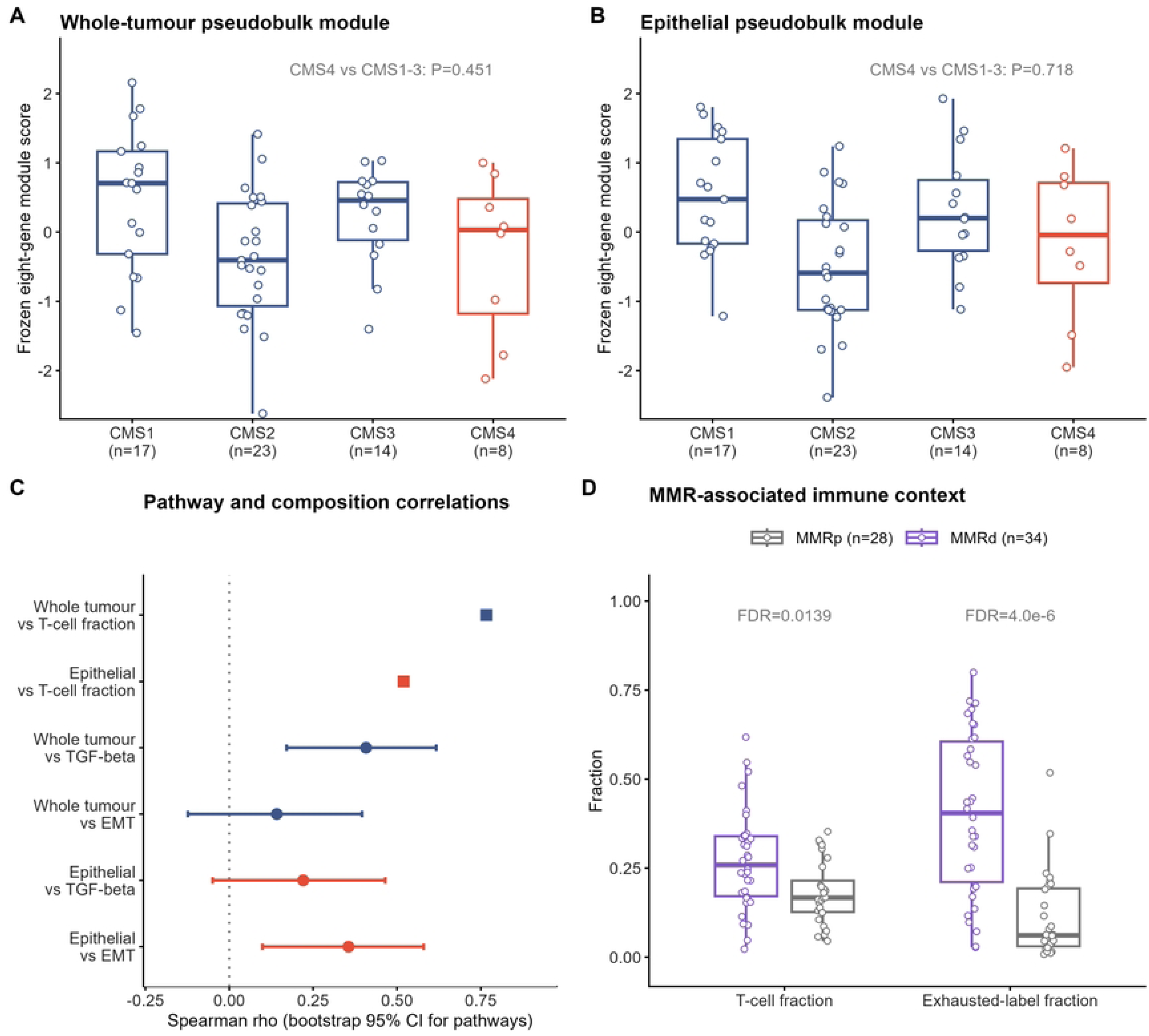
Patient-level recalibration of the GSE178341 RNA module. (A,B) Frozen eight-gene module in whole-tumour and epithelial pseudobulk by official epithelial-pseudobulk CMScaller call; CMS4 was not enriched. (C) Patient-level pathway and T-cell-composition correlations; pathway points include bootstrap 95% CIs. (D) T-cell and author-annotated exhausted-cell fractions by MMR status with BH-FDR values. Sixty-two patients (34 MMRd and 28 MMRp) were analysed. These panels describe the module’s molecular and immune context.

Whole-tumour scores correlated with TGF-beta activity (ρ=0.408; 95% CI 0.171–0.617; P=0.00100; q=0.00401), whereas epithelial scores correlated with EMT (ρ=0.356; 95% CI 0.099–0.580; P=0.00456; q=0.00913). Whole-tumour EMT and epithelial TGF-beta associations were weaker (Fig 3C). The contextual implementation also gave a positive epithelial-score association with T-cell fraction (ρ=0.520; P=1.49×10^−5^), illustrating that compartment restriction alone did not eliminate associations with the surrounding immune context.

MMRd tumours had higher median T-cell fractions than MMRp tumours (0.259 versus 0.167; P=0.00594; q=0.0139) and higher author-annotated exhausted T-cell fractions (0.405 versus 0.061; P=5.73×10^−7^; q=4.01×10^−6^). The exhaustion-expression result was concordant (P=0.000163; q=0.000572; Fig 3D).

### No survival association was detected for the specified module

In GSE39582, the hazard ratio per score SD for relapse-free survival was 0.942 (95% CI 0.796– 1.114; P=0.485) before adjustment and 1.004 (0.843–1.195; P=0.968) after adjustment for age, categorical stage, MMR and chemotherapy (Fig 4A). Models included 519 and 518 patients, respectively, with 145 events. Module Schoenfeld-residual P values were 0.590 and 0.664; the adjusted global P was 0.939.

**Fig 4.**
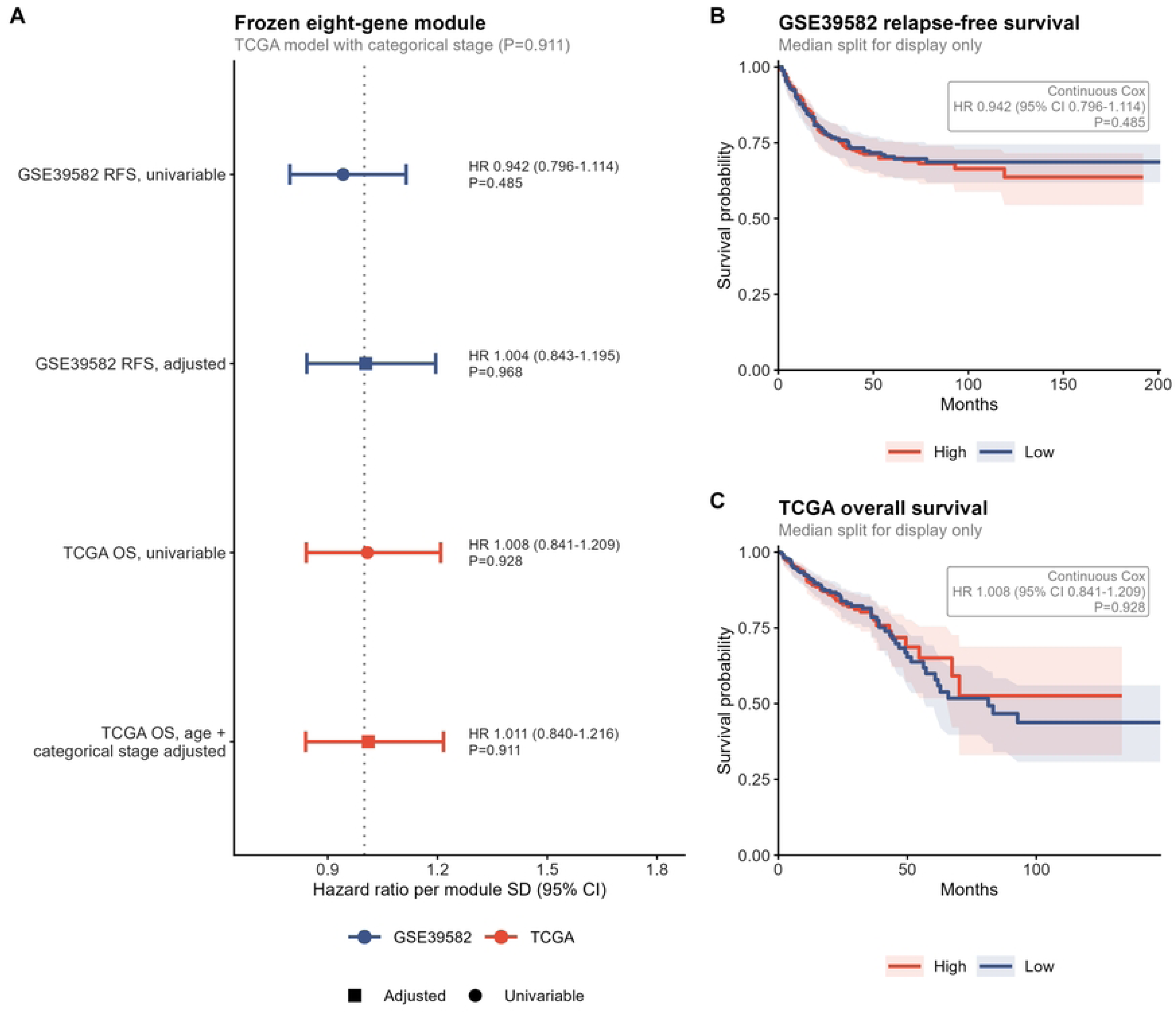
Survival analysis of the frozen eight-gene module. (A) HRs per module SD with 95% CIs and P values for GSE39582 RFS and TCGA overall survival, univariable and adjusted models. Analytic sizes were 519/518 GSE39582 cases with 145 events and 566/554 TCGA cases with 115/110 deaths. The TCGA adjusted model coded stage categorically. All module Schoenfeld-residual P values were ≥0.590. (B,C) Median-split Kaplan–Meier curves for display only; inset labels report the corresponding continuous-score univariable Cox HR, 95% CI and P value, which remain the inferential estimates.

For TCGA overall survival, the univariable estimate was 1.008 (95% CI 0.841–1.209; P=0.928; 566 patients, 115 deaths). The age- and categorical-stage-adjusted estimate was 1.011 (0.840– 1.216; P=0.911; 554 patients, 110 deaths). Corresponding module proportional-hazards P values were 0.828 and 0.873, with adjusted global P=0.701. Median-split curves are descriptive (Fig 4B,C). These intervals remain compatible with modest associations in either direction. Model estimates and proportional-hazards diagnostics are provided in S15 Table.

### Cross-platform fitness profiles of *SCD* and comparator genes

*SCD* and *EGFR* had the most negative organoid median LFCs among the 34 covered panel genes (−1.145 and −1.198), with official depletion calls in 83.5% and 82.4% of models (S1 Fig). No panel-wide MSI–MSS difference survived BH correction. Across platforms, the 34-gene median-effect correlation was modest (ρ=0.351; P=0.0418) and sensitive to removing *CD274* (ρ=0.299; P=0.0912; S2 Fig; S16 Table). *SCD* replicated as a CRC fitness dependency, with median Chronos effect −1.164 and 38/63 lines below −1; its genome-wide rank was 699/18,531.

Exploratory comparator medians in organoids were −0.089 for *CH25H*, −0.115 for *ATF3* and 0.115 for *FADS2*. Their CRC Chronos medians were 0.060, 0.036 and −0.075, respectively. No CRC-versus-other-lineage difference for these three genes passed four-target BH correction (S3A Fig; S2 Table). Near-neutral medians do not exclude effects in selected models or roles in another cell compartment.

### Adjusted co-dependency analyses and independent replication

*SCD*–*VPS72* had adjusted organoid ρ=−0.493, P=2.59×10^−6^ and single-anchor q=0.0429; pooled four-anchor q was 0.0858. The exploratory *ATF3*–*SMYD2* association had ρ=−0.525, P=4.02×10^−7^, single-anchor q=0.00666 and pooled q=0.0266 (S4A–D Fig; S3–S6 Tables).

*CH25H* and *FADS2* had no partner passing single-anchor q<0.05. Neither selected association replicated under the adjusted DepMap criteria: *SCD*–*VPS72* gave ρ=−0.175 (95% CI −0.466 to 0.093; permutation P=0.161), and *ATF3*–*SMYD2* gave ρ=0.156 (−0.128 to 0.421; P=0.230; S3B Fig; S7 Table).

No Hallmark set passed BH-adjusted P<0.05 in the original raw or adjusted *SCD* analysis (S4E Fig; S8 Table). DNA-repair enrichment weakened after adjustment (raw NES −1.513, q=0.154; adjusted NES −1.301, q=0.491), while cholesterol homeostasis was not enriched (q=0.746 and 0.890). The exploratory four-anchor extension likewise identified no pathway with within-anchor BH-adjusted P<0.05 or pooled BH-adjusted P<0.05 (S10 Table). Minimum within-anchor adjusted P values were 0.416 for *CH25H*, 0.257 for *ATF3* and 0.221 for *FADS2*; the *SCD* comparator minimum was 0.495. These values refer to the separate, uniformly implemented Hallmark 2020-labelled analysis.

## Discussion

A strong association with immune-cell abundance can be typical of matched genes rather than distinctive to a candidate biological readout. In this study, the eight-gene module lay near the centre of the matched-module correlation distribution. The tumour-wide association recurred in an independent CRC cohort. Its lower epithelial estimate was consistent with a contribution from cellular composition, although the correlation-difference interval included zero. Rare *CEACAM5*-positive leukocyte candidates provided a separate example of measurement sensitivity: their counts depended strongly on the detection threshold and were concentrated in few patients. This calibration provides reference comparisons and patient-level uncertainty for interpreting the specified RNA readouts.

The matched-null analysis separates association strength from specificity to the selected module. Immune-lineage genes are expected to covary with immune-cell abundance, so a comparison with zero correlation answers only part of the question. Matching on expression, detection and lineage specificity provided a reference distribution for comparable genes. This calibration is conditional on the matching features and eligible gene universe. Module-level expression remained imperfectly balanced; the closest-10% sensitivity was added after that imbalance was observed. Its concordant result supports the interpretation within this design, without establishing equivalence to all possible control sets.

Compartment restriction examined whether the association persisted within annotated epithelial cells. The tumour-wide association recurred across cohorts, but the 23-patient independent cohort left substantial uncertainty about attenuation. An epithelial aggregate can contain immune-marker RNA through biological transcription, ambient RNA, doublets or annotation error. The rare-transcript analysis highlights the importance of detection thresholds: most *CEACAM5*-positive candidates had low expression, and a small threshold change substantially altered their number. Paired surface-protein and contact measurements would be needed to determine their transfer origin and estimate the RNA definition’s sensitivity and positive predictive value.

These results fit the genomic context of CRC, where immune and stromal contributions shape bulk molecular classification [4, 5] and local tissue context influences epithelial plasticity [6]. They favour separating cell abundance from cell state when interpreting mixed-marker scores. The survival analyses address a further, distinct question: clinical association of the fixed score under specified endpoints and covariate adjustments. Their intervals permit modest associations in either direction. Prognostic interpretation of membrane transfer itself will require a directly measured transfer phenotype.

The functional analyses provide secondary evidence on cancer-cell fitness. *SCD* dependency was evident in CRC organoids and cell lines, consistent with established roles of SCD1 in lipid metabolism and tumour biology [18, 19]. The LSH–EWSR1 study places SCD1 in a ferroptosis-related regulatory context [20]. Cross-platform concordance supports this fitness phenotype; its relationship to membrane acquisition remains a separate experimental question. Neither the adjusted *SCD*–*VPS72* association nor the exploratory *ATF3*–*SMYD2* association met the independent cell-line replication criteria, limiting their priority as mechanistic leads.

The *CH25H*, *ATF3* and *FADS2* extension examined whether biologically motivated anchors shared a detectable fitness-associated programme in the organoid screen. No Hallmark enrichment met the specified multiple-testing threshold, so this analysis supplied no pathway-level support for such a programme. It also provides no additional test of the cellular-composition interpretation. Enrichment of fitness-correlation rankings concerns coordinated dependencies, not direct pathway activity or membrane transfer. Experimental *ATF3*–*CH25H* findings [7, 8] remain a rationale for studies in the relevant interacting cell compartments, where effects may differ from tumour-cell fitness phenotypes.

The cell compartment also matters for interpreting therapeutic SCD1 perturbation. Myeloid *Scd1* deficiency promoted an M2-like macrophage programme and CRC growth in the mouse model studied by You and colleagues [21]. By contrast, Zhao and colleagues reported reduced M2 polarisation and improved anti-PD-1 activity with a compound acting through a C/EBPβ–SCD1 pathway [22]. Genetic deletion and a compound acting through an upstream target are different interventions, so the differing phenotypes should not be reduced to a universal effect of *SCD* inhibition. Tumour-cell fitness data alone do not predict the net effect on the immune microenvironment. The phase I MTI-301 study was registered as recruiting without posted results as of 19 October 2026 [23].

A mechanistic follow-up should measure fitness and membrane acquisition in the same donor– recipient system. Traceable donor membrane proteins, recipient lineage assignment and contact controls would define the transfer phenotype. *SCD* perturbation should be assessed alongside viability, donor-marker abundance and contact frequency, with genetic rescue to test on-target effects. Flow cytometry and time-resolved imaging could distinguish acquisition from doublets or adherent material. This design would directly test whether the replicated fitness dependency also affects membrane acquisition.

Patient-level inference, matched controls, independent-cohort analysis and complete functional association tables strengthen the interpretation of these results. The author-assembled panel is heterogeneous, and its contemporaneous selection record is incomplete. Further limitations include the absence of transfer-positive reference data, imperfect matching, the small independent cohort, sparse RNA candidates and residual admixture within annotated compartments. The functional platforms differ in model composition and assay scale. The exploratory extensions depend on the selected adjustment and gene-set collection. These constraints limit generalisation beyond the specified RNA readouts and comparison conditions.

## Conclusions

Matched-module calibration showed that the fixed eight-gene RNA score tracked T-cell abundance at a strength typical of comparable immune-lineage genes. The independent cohort reproduced the tumour-wide association, while uncertainty remained about attenuation after epithelial restriction. Rare *CEACAM5*-positive leukocyte candidates were sparse and threshold-sensitive. *SCD* dependency reproduced as a separate cancer-cell fitness phenotype. Together, these findings establish an empirical basis for interpreting the tested readouts through matched comparisons, compartment-resolved analysis and patient-level uncertainty. Direct protein-acquisition measurements are needed to connect these RNA readouts to membrane transfer.

## Data availability

Organoid CRISPR data are provided with the tumour-derived organoid biobank [15]. DepMap Public 26Q1 is available from the DepMap portal [16]. TCGA-COAD/READ data were obtained from cBioPortal study coadread_tcga_pan_can_atlas_2018 [14, 24]. GSE39582 [25], GSE146771 [26], GSE178341 [27] and GSE132465 [28] are available from the NCBI Gene Expression Omnibus; these accession numbers identify the source datasets. Gene-set collections should be obtained from their providers under the applicable terms; neither Hallmark GMT is redistributed. Code, processed outputs and figure source data, including the four-anchor co-dependency and exploratory Hallmark analyses, are available at https://github.com/wangjinglin0905-gif/trogocytosis-crc-reproducibility, release v1.2.0 [29], archived at https://doi.org/10.5281/zenodo.23183934 (series DOI https://doi.org/10.5281/zenodo.22239611) [30]. S1–S16 Tables accompany this article. S11–S16 Tables reorganise existing outputs for this submission; they do not represent new analyses or identically named files in the archived release. S1 Appendix identifies additional archived outputs underlying the figures. Figure 1 has been reformatted for this submission without changing its source data. Earlier releases are retained for provenance. Third-party raw matrices are not redistributed. Code is licensed under MIT; original documentation, derived tables and figures are licensed under CC BY 4.0.

## Author contributions

JW: Conceptualization, Methodology, Investigation, Data curation, Project administration, Writing – original draft, Writing – review & editing. PS: Investigation, Data curation, Validation, Visualization, Writing – review & editing. LX: Investigation, Data curation, Validation, Visualization, Writing – review & editing. YW: Investigation, Data curation, Validation, Visualization, Writing – review & editing. WZ: Investigation, Data curation, Validation, Visualization, Writing – review & editing. YZ: Funding acquisition, Investigation, Data curation, Validation, Visualization, Writing – review & editing. YT: Investigation, Data curation, Validation, Visualization, Writing – review & editing. JY: Conceptualization, Funding acquisition, Project administration, Supervision, Writing – review & editing.

## Acknowledgments

The authors acknowledge the original data generators and public repositories.

## Supporting information

S1 Appendix. Supplementary methods and table guide. Analysis definitions, statistical implementation details, and a readable 39-gene evidence annotation (Table A) with its source bibliography.

S1 Fig. Supporting CRC organoid dependencies and subgroup comparisons. Median LFCs for the 34 covered panel genes in 85 models, *SCD* by CMS, and *SCD*/*EGFR* by microsatellite status are shown. Depletion percentages use study-provided calls, not LFC<−1. MSI–MSS q values correct across 34 genes. The endpoint is cancer-cell fitness.

S2 Fig. Supporting cross-platform functional comparisons. (A) Median-effect rank correlations for 34 shared genes and a 33-gene sensitivity excluding *CD274*. (B) Official organoid depletion rates and DepMap Chronos<−1 rates are different descriptive criteria. (C) Original *SCD*/*EGFR* CRC-minus-other-lineage median differences with bootstrap intervals and Mann–Whitney P values. The *EGFR* interval includes zero despite a nominal P<0.05; the rank-distribution test and median-difference interval summarise different estimands.

S3 Fig. Exploratory lineage comparisons and cross-platform association checks. (A) CRC-minus-non-CRC median Chronos effects in 63 and 1,145 cell lines, with 10,000-resample bootstrap intervals and BH correction across four targets. Intervals use seed 42; Monte Carlo endpoints need not exactly match the original *SCD* interval in S2 Fig. (B) Adjusted *SCD*–*VPS72* and *ATF3*–*SMYD2* associations in 63 CRC lines, using 1,179 common-essential genes defined across all cell lines. Intervals use 10,000 line bootstraps with reranking/refitting; P values use 10,000 residual permutations. Neither pair meets the replication criteria.

S4 Fig. Adjusted organoid associations and original *SCD* pathway analysis. (A–D) Each anchor is excluded from its 16,561-partner family; the ordinate is −log10 of the single-anchor BH q value. Three covariates imply df=80 for the approximate partial-correlation test. Labels additionally give pooled four-anchor q values. *SCD*–*VPS72* passes the single-anchor but not the pooled 0.05 threshold; *ATF3*–*SMYD2* passes both. (E) Two biologically relevant pathways from the complete original *SCD* Hallmark analysis, not a selection of significant sets. No pathway in either full ranking has q<0.05; complete results are provided in S8 Table. The separate four-anchor exploratory Hallmark analysis is reported in S10 Table.

S1 Table. Evidence annotation and screen coverage of the 39-gene panel.

S2 Table. Four-target CRC-versus-other-lineage effects and uncertainty.

S3 Table. Complete adjusted *SCD* organoid co-dependency results.

S4 Table. Complete adjusted *CH25H* organoid co-dependency results.

S5 Table. Complete adjusted *ATF3* organoid co-dependency results.

S6 Table. Complete adjusted *FADS2* organoid co-dependency results.

S7 Table. Independent DepMap replication checks.

S8 Table. Complete original *SCD* Hallmark analysis.

S9 Table. Pooled four-anchor organoid associations and BH correction. The full 66,244-row table is supplied as a gzip-compressed CSV.

S10 Table. Complete exploratory four-anchor Hallmark analysis and both BH correction families.

S11 Table. GSE178341 matched-null calibration and matching balance. Empirical calibration of the frozen eight-gene module against 10,000 expression-, detection- and lineage-matched random modules at k=25, 50 and 100, with upper-tail, lower-tail and equal-tail two-sided empirical probabilities, and the three frozen-module means relative to their matched-null distributions.

S12 Table. GSE178341 patient-level module scores and T-cell fractions. Complete-gene CPM and archived scores for the 62 tumour patients.

S13 Table. GSE132465 compartment replication. Cohort-level correlation, permutation and bootstrap summaries for 23 patients, together with the tumour-wide and epithelial patient-level scores.

S14 Table. GSE146771 *CEACAM5*-positive leukocyte candidates. Tissue-level detection rates with Wilson 95% CIs and patient-, tissue- and lineage-level candidate counts across the author-annotated leukocyte atlas.

S15 Table. GSE39582 and TCGA survival models with proportional-hazards diagnostics. Hazard ratios per score SD with 95% CIs for univariable and adjusted models, and the TCGA cox.zph term and joint GLOBAL tests.

S16 Table. Cross-platform organoid–DepMap panel effects and rank correlation. Organoid median knockout LFC and DepMap 26Q1 CRC Chronos medians with interquartile values for the 34 shared panel genes, and Spearman rank correlations for the 34-gene and 33-gene (*CD274*-excluded) definitions.

